# MRE11A: a novel negative regulator of human DNA mismatch repair

**DOI:** 10.1101/2023.03.15.532782

**Authors:** Demin Du, Yueyan Yang, Guanxiong Wang, Liying Chen, Xiaowei Guan, Lene Juel Rasmussen, Dekang Liu

## Abstract

DNA mismatch repair (MMR) is a highly conserved pathway that corrects DNA replication errors. Although well characterized, MMR factors remain to be identified. As a 3’-5’ exonuclease and endonuclease, meiotic recombination 11 homolog A (MRE11A) is implicated in multiple DNA repair pathways. However, the role of MRE11A in MMR is unclear. Here, we show that MRE11A deficiency increased the sensitivity of HeLa cells to N-methyl-N’ nitro-N nitrosoguanidine (MNNG) treatment, implying a potential role of MRE11 in MMR. Moreover, we found MRE11A was largely recruited to chromatin and negatively regulated the DNA damage signals within the first cell cycle after MNNG treatment. We also showed that knockdown of MRE11A increased, while overexpressing MRE11A decreased, MMR activity in HeLa cells, suggesting that MRE11A negatively regulates MMR activity. Furthermore, we show that the recruitment of MRE11A to chromatin requires MLH1 and that MRE11A competes with PMS2 for binding to MLH1. This decreases PMS2 levels in whole cell and on chromatin, and consequently comprises MMR activity. Collectively, our findings reveal that MRE11A is a negative regulator of human MMR.

## INTRODUCTION

High fidelity of DNA replication is critical to maintain genomic integrity during cell proliferation. DNA mismatch repair (MMR) is an important DNA repair pathway that plays critical role in DNA replication fidelity. MMR is composed of three main steps, including the mismatches recognition, mismatches removal and the DNA strand resynthesis. In human cells, MutSα (MSH2-MSH6 heterodimer) and MutSβ (MSH2-MSH3 heterodimer) are responsible for recognizing and binding to the base-base mismatches and larger insertion/deletion loops (IDLs) respectively. MutLα (MLH1-PMS2 heterodimer) is recruited by MutSα/β, forming a tetrameric complex sliding clamp to produce nicks adjacent to the mismatches, from which Exonuclease 1 (EXO1) excises the mismatch containing strand, and the leaving gap is refilled by DNA polymerase δ, and nick is re-ligated by DNA ligase I (Fishel, 2015; Liu *et al*, 2017). Loss of MMR led to hyper-mutational phenotypes in human cells that contributed significantly to early-onset of various types of cancers (Lynch *et al*, 2015; Stelloo *et al*, 2016). In clinic, MMR status is widely used as biomarker for diagnosis and medication choice for cancer treatment. For instances, MMR-deficient tumors exhibit resistance to chemotherapeutic drugs, such as alkylating agents (Baretti & Le, 2018; Battaglin *et al*, 2018; Irving & Hall, 2002; Leelatian *et al*, 2021; Pors & Patterson, 2005). Besides, MMR-deficient tumors are sensitive to immunotherapy, *i*.*e*., anti-PD-1 or PD-L1 treatment (Majidpoor & Mortezaee, 2021; Nebot-Bral *et al*, 2019; Oliveira *et al*, 2019). As such, evaluating MMR activity in tumors is critical for early diagnosis and treatment of cancer. For assessing MMR status, immunostaining of known MMR factors or microsatellite instability (MSI) test are mostly used in clinic (Pena-Diaz & Rasmussen, 2016). However, a large proportion of MSI cancers cannot be assessed by alternations of known MMR proteins (Baretti & Le, 2018; Shia, 2008), indicating that there are still unknown MMR factors that remain to be identified. Given that EXO1 deficiency only causes milder MMR-deficient phenotypes compared to inactivation of MSH2 or MLH1 (Jagmohan-Changur *et al*, 2003; Szankasi & Smith, 1995; Wei *et al*, 2003), we initiated a screen aiming to identify novel exo/endonucleases in MMR other than EXO1 (Song *et al*, 2022). We found that knockdown of meiotic recombination 11 homolog A (MRE11A) enhanced the sensitivity of cells to the alkylating drug N-methyl-N’nitro-N-nitrosoguanidine (MNNG).

MRE11A protein possesses 3’-5’ nuclease and endonuclease activity. In cells, MRE11A assembles with two other partners RAD50 and NBS1, forming the MRN complex, which serves as the early sensor and processor of DNA double strand breaks (DSBs) and is engaged in many chromosome metabolic processes, such as DNA homologous recombination, telomere metabolism, DNA replication forks processing and meiosis (Carney *et al*, 1998; Farah *et al*, 2009; Langerak *et al*, 2011; Paull & Gellert, 1998; Syed & Tainer, 2018). Besides, MRE11A also participates in break induced repair/replication (BIR) and interstrand crosslink (ICL) repair (Furuta *et al*, 2003; Kramara *et al*, 2018; Stingele *et al*, 2017). Furthermore, it interacts with MLH1, indicating a role for MRE11A in MMR (Mirzoeva *et al*, 2006; Zhao *et al*, 2008). Previously it was reported that MRE11A-deficient cells show reduced MMR activity on 3’ nicked DNA substrates *in vitro* (Vo *et al*, 2005), whereas others argued that MMR deficiency also affects MRE11A expression or functions (Franchitto *et al*, 2003; Gaymes *et al*, 2013; Giannini *et al*, 2002; Ham *et al*, 2006; Wen *et al*, 2008). One later study using murine cells showed that inactivation of MRE11A had no influence on MMR activity (Desai & Gerson, 2014). These divergent findings prompted us to further characterize the role of MRE11A in MMR. Here, we show that inactivation of MRE11A increases cells’ sensitivity to MNNG, and demonstrate that MRE11A negatively regulates MMR activity through competing with PMS2 for binding to MLH1.

## RESULTS

### MRE11A knockdown increases the sensitivity of HeLa cells to MNNG

In the absence of MGMT, MNNG can generate ^O(6)Me^G lesions, and the ^O(6)Me^G:T mis-pairs are recognized by MutSα during genomic replication. If O^6^MeGs are located on mother-strand, the MMR machinery may enter “futile cycle”, in which the thymine is repeatedly excised and mis-incorporated opposite O^6^MeG. Thus, nicks or gaps may persist, and can be converted into a replication fork collapse in the second round of genomic replication, which then activates a G2 checkpoint and subsequent cell cycle arrest (Mojas *et al*, 2007; Quiros *et al*, 2009; Song *et al*., 2022). Therefore, higher MMR activity correlates with greater sensitivity to MNNG in MGMT inactivated cells. We observed that MRE11A-depleted cells demonstrate increased sensitivity to MNNG in our screening assay (Song *et al*., 2022). To validate MRE11A as a positive hit, two different siRNA targeting MRE11A in HeLa cells, siMRE11A-1 and siMRE11A-2, were used. The siNC and siMLH1 were used as controls in both assays. The viability of the MRE11A depleted cells were measured 72 hours (after approximately 3 cell cycles) after MNNG treatment (O^6^-Benzylguanine was added 1 hour before to inactivate MGMT). As shown in **Figure 1A**, MRE11A deficient cells exhibited abnormal morphology and less survival compared to siNC control after MNNG treatment at the concentration as low as 200nM (*P*=0.002, 0.002). In parallel, clongenic assay showed that MRE11A knockdown reduced the percentage of surviving cell colonies to siNC control after MNNG treatment at the concentrations of 50nM (*P*=0.0012,0.016), 100nM (*P*=0.019, 0.087), and 150nM (*P*=0.071, 0.031) **(Figure 1B)**. While, there was no significant differences in the growth rate and protein expression levels of MSH2 and MLH1 after MNNG treatment between MRE11A knockdown and siNC control cells **(Figure 1C, D)**.

**Figure 1.**
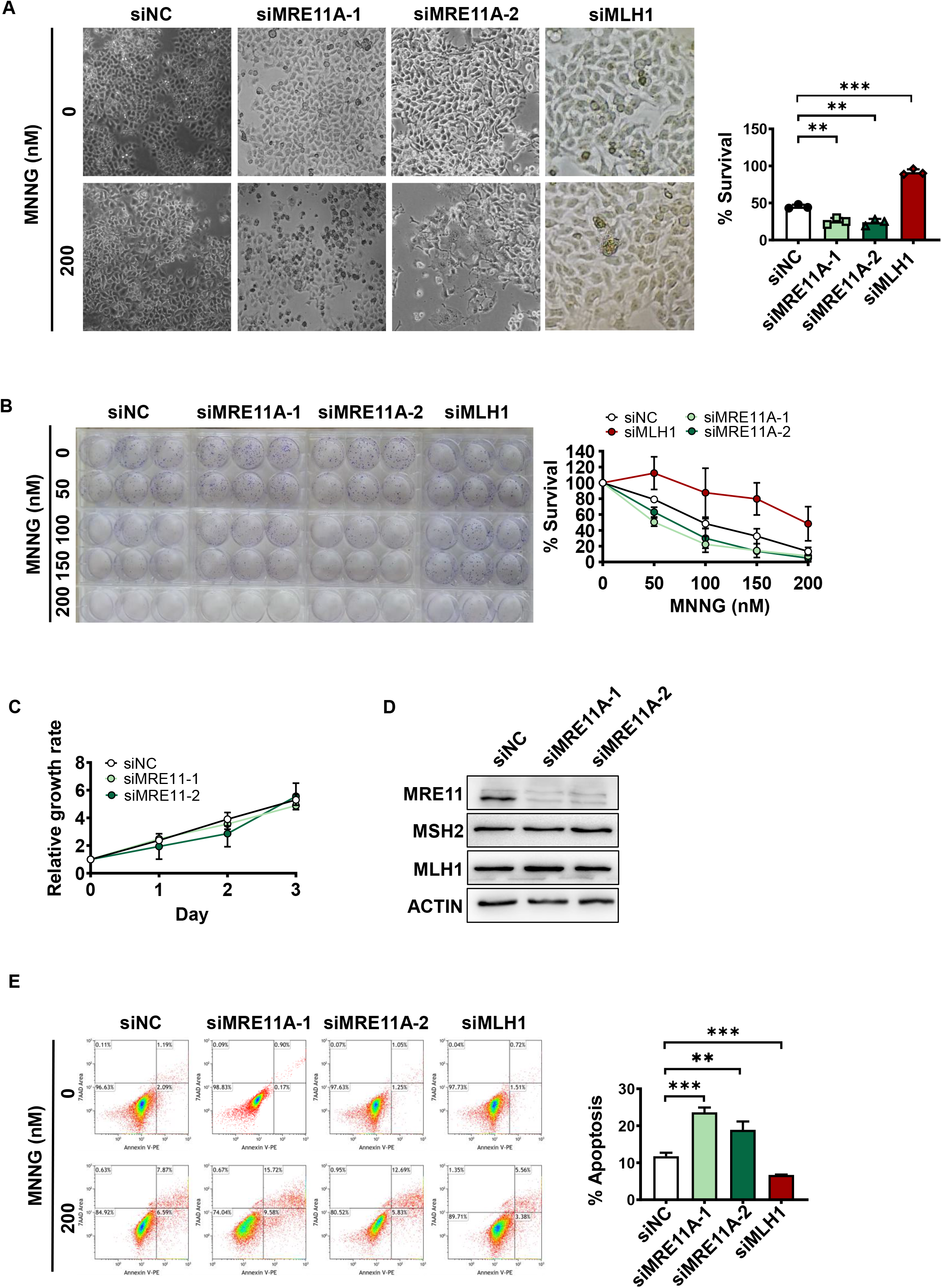
MRE11A deficiency sensitizes cells to MNNG treatment. **(A)** Hela cells were transfected with two different siRNAs targeting MRE11A and exhibited abnormal morphology and less survival 72 hours after 200nM MNNG treatment. The survival rate was the percentage of survival cells to the parallel cells treated only with O6-Benzylguaine and DMSO in each group, and cells with MLH1 deficiency was set as positive control. **(B)** MRE11A knockdown cells were treated with MNNG and triplicated seeded in 6 well plates, and after approx. 2 weeks, cells were stained by crystal violet and the clones with ≥200 cells were counted. The survival rate was the percentage of survival clones to the parallel wells treated only with DMSO in each group, and cells with MLH1 deficiency was set as positive control. **(C)** The growing rates of control cells and MRE11A knockdown cells were measured in 96 well plates with CCK8 reagents. **(D)** Western blotting picture exhibited no significant changes in proteins levels of MSH2 and MLH1 in MRE11A knockdown cells. **(E)** Representative flow cytometry pictures of scatter plots of PI *vs* Annexin V staining of the siNC control, MRE11A and MLH1 knockdown cells 72 hours after 200nM MNNG treatment. The right graph showed the statistical analysis of the left flow cytometry data, quantification and comparison of the proportions of apoptosis cells in each group. The %apoptosis was calculated as the %apoptosis of cells with 200nM MNNG minus that with only DMOS treatment. All data were analyzed with unpaired two-tailed Student’s t test. Data were shown as mean ± SD, n = 3, * *P* < 0.05, ** *P* < 0.01, *** *P* < 0.001.

When cells are challenged with MNNG, the MMR machinery may activate cell apoptosis pathways after two cell cycles (Quiros *et al*., 2009). We found that knockdown of MRE11A increased apoptosis compared to siNC controls (*P*<0.001, =0.0069), while knockdown of MLH1 decreased apoptosis as expected (*P<*0.001) **(Figure 1E)**. Together, these results suggest that MRE11A depletion sensitizes cells to alkylation damage (such as MNNG), implying the possible regulatory role for MRE11A in MMR.

### MRE11A depletion enhances MNNG-induced DNA damage signals within one round of cell cycle

MNNG-induced cell death requires MMR activity to introduce replication fork collapse in two rounds of cell cycle (Mojas *et al*., 2007; Quiros *et al*., 2009). MRE11A also participates in the repair of replication fork collapse (Furuta *et al*., 2003; Kramara *et al*., 2018). Therefore, in order to avoid the interference of the role of MRE11A in replication fork collapse repair and investigated the specific role of MRE11A in MMR, we treated HeLa cells with MNNG for 12 hours, a period within one round of cell cycle **(Figure 1D**). As shown in **Figure 2A**, MRE11A is already recruited to chromatin 12 hours after MNNG treatment (*P*=0.047), suggesting a role for MRE11A in processing alkylation damage together with MMR. The phosphorylation level of CHK1 and number of 53BP1 foci in cells in G1 phase were measured to evaluate extent of DNA damages caused by MMR processing of alkylation damage in the first cell cycle after MNNG treatment (Gupta *et al*, 2018; Lukas *et al*, 2011; Mojas *et al*., 2007; Wang & Qin, 2003; Yoshioka *et al*, 2006). The results showed that MRE11A deficiency increased both CHK1 levels and the number of 53BP1 foci after MNNG treatment compared to controls (*P*=0.041, 0.011 and 0.0040, <0.001) **(Figure 2B, C)**. To consolidate our findings, we overexpressed MRE11A in HeLa cells and found decreases in CHK1 phosphorylation level (*P*=0.0015) and 53BP1 foci numbers (*P*= 0.0053) compared to controls **(Figure S1)**. These results suggest that MRE11A prevents MNNG-induced DNA damage potentially through interfering MMR activity.

**Figure 2.**
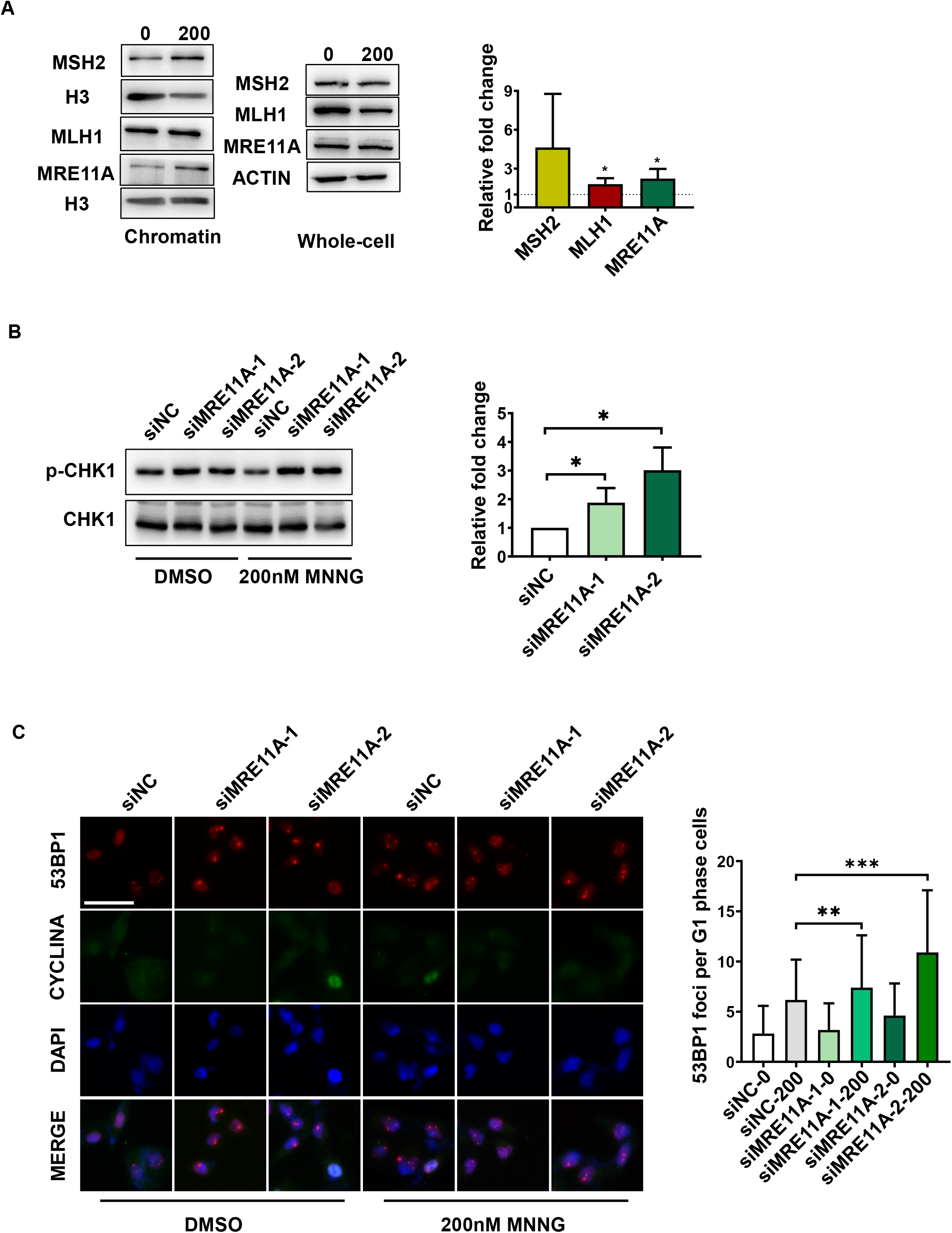
MRE11A deficiency increases DNA damage signals 12h after MNNG treatment. **(A)** The representative western blotting pictures of the chromatin binding and whole cell MSH2, MLH1 and MRE11 proteins 12 hours after DMSO or 200nM MNNG treatment of Hela cells. The level of histone H3 was set as inner control. Right graph was the quantification of fold change of ratio of the chromatin binding to the whole cell proteins of MSH2, MLH1 and MRE11A after exposure to 200nM MNNG. **(B)** The representative western blotting of the phosphorylation levels of CHK1 12 hours after DMSO or 200nM MNNG treatment. The alternation of phosphorylation level was calculated as p-CHK1 level normalized by total CHK1 protein after 200nM MNNG treatment minus that with only DMSO treatment. Right graph showed the quantification of proteins level changes relative to siNC. **(C)** Representative Immunofluorescent pictures of the 53BP1 foci in G1 phase 12 hours after DMSO or 200nM MNNG treatment. Right graph showed the quantification of the number of 53BP1 foci per cell in G1 phase (CYCLINA-). At least 250 cells were counted for each group. Data are shown as mean ± SD, * *P* < 0.05, ** *P* < 0.01, *** *P* < 0.001, using unpaired two-tailed Student’s t test.

### MRE11A negatively regulates MMR activity

To explore the impact of MRE11A on MMR activity, we employed a GFP-heteroduplex assay. This assay relies on the detection of GFP expression as a marker of successful heteroduplex repair. Additionally, co-transfection of mcherry plasmids was implemented to normalize transfection efficiency across all assays (Zhou *et al*, 2009). We found that MRE11A knockdown increased MMR activity (*P*=0.0015, 0.0087, **Figure 3A**), and its overexpression decreased MMR activity (*P*=0.0067, **Figure 3B**), in accordance with the above results that MRE11A can negatively regulate MMR-induced DNA damage signals as well as cell survival after MNNG treatment.

**Figure 3.**
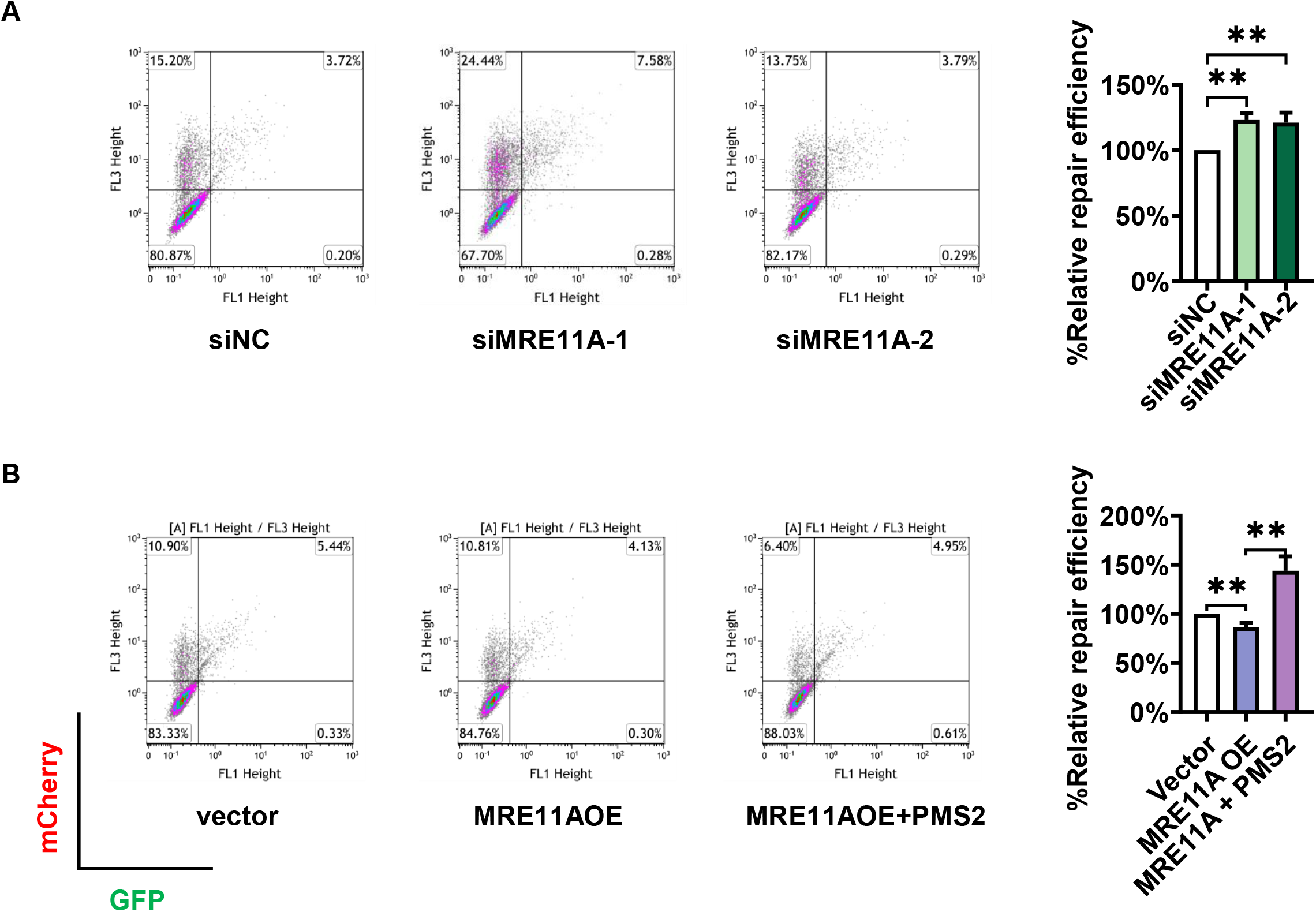
MRE11A negatively regulates MMR activity in Hela cells. Left pictures represented the scatter plots of cells co-transfected with GFP-heteroduplex and mcherry plasmids described in materials and methods. The x-axis and y-axis represented the signal intensities of GFP and mcherry respectively. The MMR repair efficiency was calculated as the ratio of the number of GFP positive cells to mcherry positive cells, and the quantification results relative to siNC or empty vector controls were in the right graphs. Data are shown as mean ± SD, * *P* < 0.05, ** *P* < 0.01, *** *P* < 0.001, using unpaired two-tailed Student’s t test.

Microsatellite Instability (MSI) is an indirect indicator for MMR deficiency (Saha *et al*, 2005). We established stable cell lines of MRE11A knockdown or overexpression, duplicated single cell approximately 30 times (around 30 days), and harvested genomic DNA to access the insertion/deletion mutations of microsatellite markers, BAT25, BAT26, MONO27, NR21 and NR24. Surprisingly, we did not detect any changes of these markers **(Figure S2A)**, while MSI was detected in the positive control of HEK293T cells (MLH1-deficient) **(Figure S2B)**. These results suggest that MRE11A negatively regulates MMR activity, but not to an extent that can affect the stability of microsatellites within approximately 30 genome duplications.

### MRE11A is recruited to chromatin by MMR protein

Since MRE11A can regulate MMR activity, we sought to investigate whether it interacted with the MMR machinery. We performed chromatin immuno-precipitation with an MSH2 antibody and observed that without MNNG treatment, MSH6 was recruited with MSH2 on chromatin as expected, while very low levels of MLH1 and MRE11A were co-precipitated **(Figure 4A)**. 12 hours after MNNG treatment, we found co-recruitment of MRE11A and MLH1 on chromatin with MSH2, implying an interaction between MRE11A and MLH1 **(Figure 4A)**. Moreover, MSH2 deficiency downregulated, though not significantly (*P*=0.07), MRE11A level on chromatin in untreated cells, while MLH1 deficiency significantly decreased MRE11A on chromatin (*P*<0.001) **(Figure 4B)**. Meanwhile, we found that MRE11A deficiency did not influence the levels of chromatin bound MLH1 and MSH2 **(Figure 4C)**. These results support previous findings that MRE11A co-precipitated with MLH1 after MNNG treatment (Vo *et al*., 2005; Zhao *et al*., 2008), and collectively suggest that MRE11A participates in the MMR pathway through interaction with MLH1.

**Figure 4.**
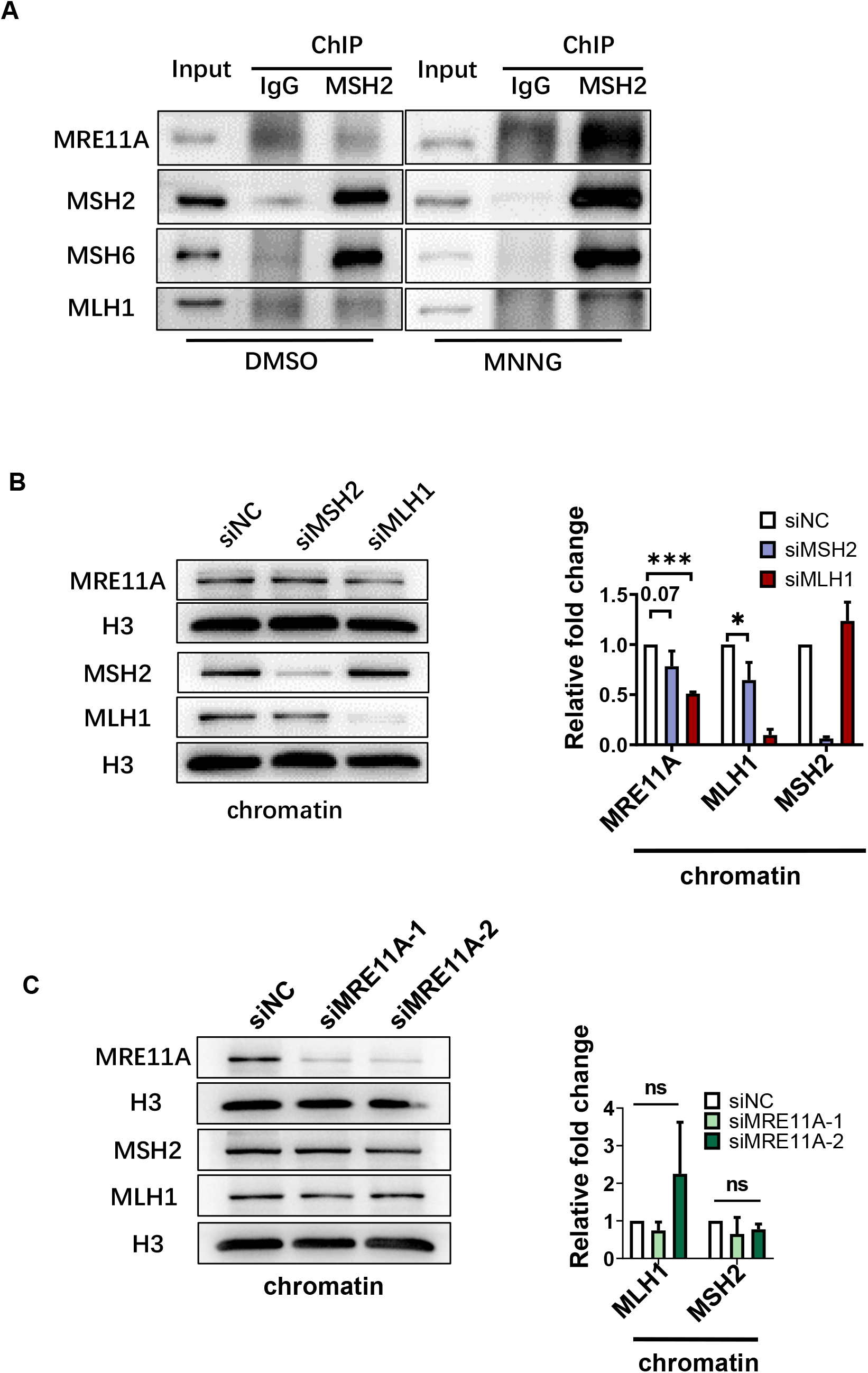
MRE11A is recruited to chromatin by MMR protein. **(A)** Representative Western blotting of the chromatin proteins co-precipitated with MSH2 12 hours after DMSO or 200nM MNNG treatment. **(B, C)** Representative Western blotting of the chromatin binding MSH2, MLH1 and MRE11A proteins after knockdown of MSH2, MLH1 and MRE11A independently. The right graphs were the quantification of the western blotting bands relative to siNC controls. Data are shown as mean ± SD, * *P* < 0.05, ** *P* < 0.01, *** *P* < 0.001, using unpaired two-tailed Student’s t test.

### MRE11A negatively regulates PMS2 protein levels

The interaction domain of MLH1 with MRE11A is overlapping that with PMS2, which is situated at the C-terminal of MLH1 around amino acids 495-756AA. Additionally, mutations in K618, K616 or L574 of MLH1 disrupt MLH1 interactions with MRE11A and PMS2 simultaneously (Guerrette *et al*, 1999; Plotz *et al*, 2003; Vo *et al*., 2005; Zhao *et al*., 2008). Hence, it was conceivable that MRE11A may compete with PMS2 for binding to MLH1, which downregulates the level of MLH1**-**PMS2 on chromatin and consequently interfere with MMR activity. To verify our hypothesis, we examined the PMS2 protein level and found that it was decreased (*P*=0.0032) when MRE11A was overexpressed and increased (*P*=0.0086, 0.0050) when MRE11 was depleted, while the mRNA levels remained unchanged **(Figure 5A, B)**, which could be explained by the fact that PMS2 protein is unstable in absence of MLH1 binding (Bellizzi & Frankel, 2009; Kosinski *et al*, 2010). As expected, PMS2 levels on chromatin were decreased (*P*<0.001) when MRE11A was overexpressed and increased when MRE11A was depleted **(Figure 5C, D)**, while MLH1 levels remained unchanged **(Figure 4C and Figure 5A, B, D)**. Next, we found that PMS2 overexpression can restore the decreased MMR activity (*P*=0.0029) caused by MRE11A overexpression **(Figure 3B)**, indicating MRE11A expression interfered with MMR activity via downregulating PMS2 level. Furthermore, we found that overexpression of MRE11A peptide 452-634AA, the binding motif with MLH1 (Vo *et al*., 2005; Zhao *et al*., 2008), can decrease PMS2 protein level in whole cell extracts (*P*=0.0013) as well as on chromatin (*P*=0.0046) **(Figure 5B, D)**. We also find that overexpression of MRE11A peptide 452-634AA downregulated MMR activity (*P*<0.001), which can be rescued by PMS2 overexpression (*P*<0.001) **(Figure 5E)**.

**Figure 5.**
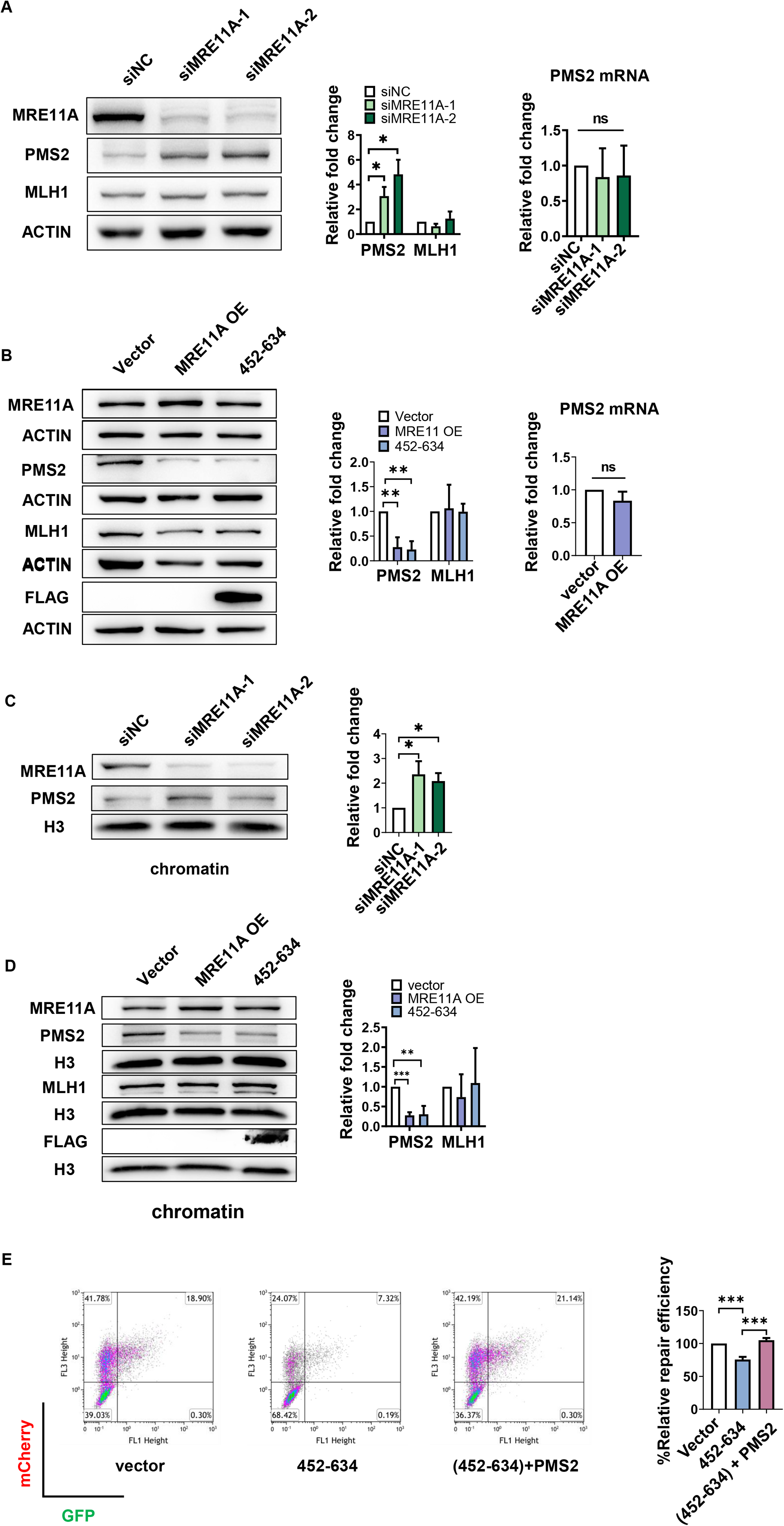
MRE11A negatively regulates PMS2 levels. Representative western blotting of the indicated proteins levels in whole cells **(A, B)** or on chromatin **(C, D)** with MRE11A knockdown/overexpression or expression of flag-tagged 452-634AA of MRE11A. The right graphs were the quantification of the western blotting intensities relative to siNC or empty vector controls. The changes of mRNA levels of PMS2 after MRE11A knockdown **(A)** or overexpression **(B)** were quantified using qPCR. The representative scatter plots of flow cytometry analysis of cells expressing GFP or mcherry, reflecting MMR repair efficiencies of cells expressing 452-634AA of MRE11A with/wo PMS2 overexpression **(E)**. The MMR repair efficiency was calculated as the ratio of the number of GFP positive cells to mcherry positive cells, and the quantification results relative to siNC or empty vector controls were in the right graphs. Data are shown as mean ± SD, * *P* < 0.05, ** *P* < 0.01, *** *P* < 0.001, using unpaired two-tailed Student’s t test.

To test if MRE11A can regulate PMS2 levels in other cell lines, MRE11A was depleted in SH-SY5Y (neuroblastoma), MDA-MB231 (breast cancer), A549 (lung cancer), HepG2 (liver carcinoma), and U87mg (glioblastoma) cells. The results showed that MRE11A knockdown in these cell lines also upregulate PMS2 levels **(Figure 6)**, verifying that our results were not specific for HeLa cells. Taking together, these results show that MRE11A interferes MMR activity by competing with PMS2 for binding to MLH1.

**Figure 6.**
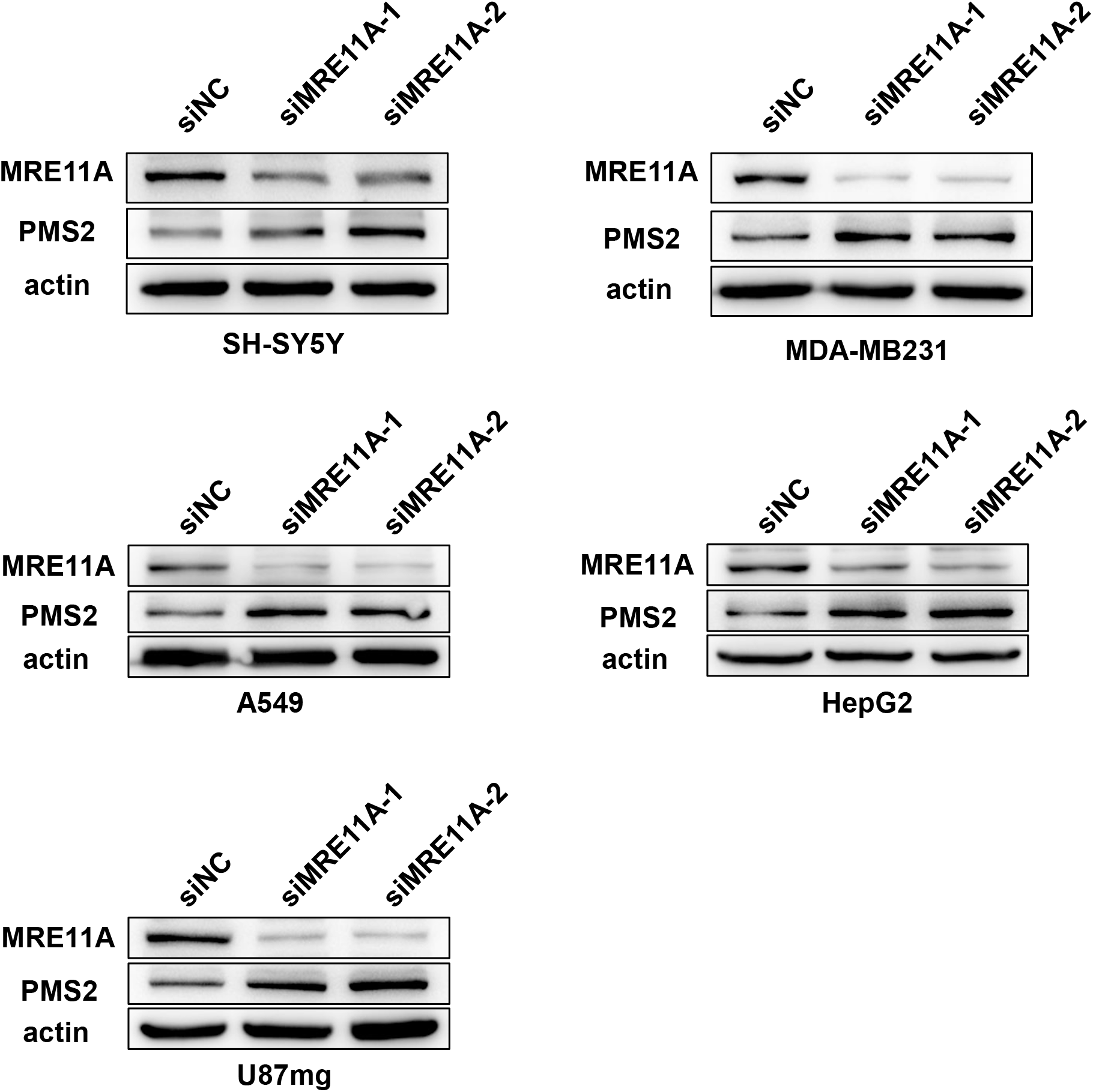
MRE11A knockdown increases PMS2 levels in various cell lines. Representative western blotting of the PMS2 protein levels in whole cell lysis after MRE11A knockdown in indicated cell lines.

## DISCUSSION

Mismatch repair is a highly conserved physiological process from prokaryote to eukaryote and plays irreplaceable roles in maintenance of genomic integrity. In prokaryote, multiple helicases and nucleases, such as Exo1, ExoVII, ExoX and RecJ, contribute to the removal of mispaired bases in the newly synthesized strand. In contrast, Exonuclease 1 (EXO1) was recognized as the only exonuclease involved in eukaryotic MMR (Kadyrov *et al*, 2006). Nevertheless, mounting investigations illustrated that depletion of EXO1 in yeast or human cells only led to mild downregulation of MMR activity, indicating alternative factors in the excision step (Jagmohan-Changur *et al*., 2003; Kadyrov *et al*, 2009; Thompson *et al*, 2004; Tishkoff *et al*, 1997; Wei *et al*., 2003). To date, reports showed that the strand displacement activity of DNA Polδ, WRN helicase, FAN1 or synergetic effect of multiple nucleases may act secondary to EXO1 in MMR (Kadyrov *et al*., 2009; Kratz *et al*, 2021; Picco *et al*, 2021). Previously, we performed high throughput screening for novel MMR nucleases or helicases using cellular sensitivity to low dosage MNNG (belongs to the SN1 alkylation reagent) as a read out for MMR activity (Song *et al*., 2022). Among the candidates, we found knockdown of MRE11A induced increased sensitivity of HeLa cell to MNNG, implying that it may negatively regulate MMR activity. This finding was unexpected, since Her’s group previously reported that MRE11A-deficient cells showed microsatellite instability on artificial substrates and comprised 3’ nick directed MMR activity *in vitro* (Vo *et al*., 2005). Given that MRE11A possesses 3’-5’ exonuclease activity and two reports demonstrated that MRE11A can interact with MLH1, it was naturally to propose MRE11A as an alternative nuclease in MMR (Vo *et al*., 2005; Zhao *et al*., 2008) (Giannini *et al*., 2002; Wu *et al*, 2011). However, two reports have shown that inactivation of MRE11A in eukaryotes did not affect the MMR activity (Desai & Gerson, 2014; English, 2007). Considering the existence of these divergent findings and the identification of MRE11A as the candidate in our screening assay, we performed an in-depth study of MRE11A to verify its role in human MMR.

Initially, to validate the screening result, we used two different siRNA to transient downregulate MRE11A level and re-confirm MRE11A deficiency increased cells’ sensitivity to MNNG with short-term and long-term clongenic assay. MRE11A may also participates in repair of replication forks collapse in the second round of cell cycle after MNNG treatment, so in order to characterize the specific role of MRE11A in MMR, we evaluated the level of CHK1 phosphorylation and the number of 53BP1 foci within the first cell cycle after MNNG treatment. We performed MMR activity assay, directly showing that MRE11A levels negatively correlated with MMR activity. We also demonstrated that MRE11A could be recruited to chromatin with MLH1 and interfered the binding of PMS2 to MLH1, leading to decreased MLH1-PMS2 levels on chromatin, by which negatively regulated MMR activity.

MRE11A plays multifaceted functions in DNA repair and metabolism, such as repair of DSBs repair and the processing of stalled or collapsed replication forks (Farah *et al*., 2009; Lamarche *et al*, 2010; Langerak *et al*., 2011; Paull & Gellert, 1998; Syed & Tainer, 2018). Although we observed increased DNA damage signals in MRE11A knockdown cells 12 hours after MNNG treatment and strong recruitment of MRE11A to chromatin by MLH1, we cannot exclude alternative roles of MRE11A in processing of DNA lesions or replication stress. Indeed, one report showed that HeLa cells encountered replication stress and retarded S phase progression in the first cell cycle after MNNG treatment (Gupta *et al*., 2018). Hence, the increased DNA damage signals may be attributed to the insufficient processing of replication stress in absence of MRE11A. However, DNA damage response to the replication stress after MNNG treatment has yet to be clarified, and until now the MMR directed repair of ^O(6)Me^G/T remains the cause for activation of DNA damage signals within the first cell cycle.

We found MRE11A negatively regulates MMR activity when performing assays that measures repair mismatches on artificial substrates in cells. To verify our results furtherly, we conducted microsatellite instability (MSI) analysis, but observe no instability of five clinically verified microsatellites. Microsatellites are repetitive sequences, commonly 1-6 bases, in the genome.

DNA polymerases are inefficient at replication of microsatellite sequences, leading to the generation of replication slippages, which are then corrected by MMR to avoid the insertion/deletion mutations. Hence, MMR-deficient cells have high probability to stochastically accumulate mutations in microsatellite sequences. Prevention of MSI seems to require almost complete MMR activity since inactivation of MSH2 and MLH1 causes high MSI (Pinol *et al*, 2005), whereas deletion of EXO1 did not necessarily cause MSI (Alam *et al*, 2003; Wei *et al*., 2003; Wu *et al*, 2001). Therefore, although MSI was thought to be the hallmark for MMR deficiency, cells with decreased MMR activity may not develop MSI within a limited number of duplications. Therefore, sensitive detection methods might be required, such as NGS based MSI testing. These results may explain why we could not detect MSI in MRE11A-deficient cells.

We and others have recently identified factors that negatively regulate MMR activity such as SLX4, CNOT6, HDAC6 and FAN1 (Goold *et al*, 2018; Guervilly *et al*, 2022; Song *et al*., 2022; Zhang *et al*, 2014). Additionally, here we report that MRE11A inhibits MMR repair by competing with PMS2 for complex formation with MLH1. The discovery of negative MMR regulators may imply the mechanism cells employ to avoid potential damaging effect brought by unbalanced MMR activity. One well-known effect of unbalanced MMR is the expansion of triplet nucleotides repeats (mutagenic effect), which may subsequently lead to the onsets of many human diseases, such as Fragile X syndrome, Huntington’s disease and Spinocerebellar ataxia etc. (Iyer *et al*, 2015). Presumably, excessive MMR activity would cause cellular hypersensitivity to alkylation damage and cells would undergo unnecessary processing of uncanonical DNA structures, causing chromatin instability. Indeed, studies have shown that overexpression of MSH3, MLH1 or PMS1 cause downregulation of MMR activity or hyper-mutational phenotypes in eukaryotes cells (Gibson *et al*, 2006; Marra *et al*, 1998; Shcherbakova *et al*, 2001). Furthermore, clinical studies indicated the correlation of overexpression of MMR proteins with elevated tumor aggressiveness or poor prognostics in various types of cancer (Albero-Gonzalez *et al*, 2019; Chakraborty *et al*, 2018; Huang *et al*, 2017; Kauffmann *et al*, 2007; Li *et al*, 2008; Liccardo *et al*, 2020; Velasco *et al*, 2002; Wagner *et al*, 2016; Wilczak *et al*, 2016). Hence, the negative regulators of MMR seem crucial to maintain a critical level of MMR to ensure genomic stability and metabolic homeostasis.

The existence of gaps or nicks on the mismatches-containing DNA strands are indispensable for correcting mismatches. In canonical MMR pathway, the intrinsic endonuclease activity of PMS2 is activated when MLH1-PMS2 heterodimer is formed, creating nicks adjacent to mismatches for the entry for exonuclease. Hence, the decrease of PMS2 protein may severely compromise the MMR activity (Kasela *et al*, 2019). Here we show that MRE11A can compete with PMS2 for binding to MLH1, consequentially interfering the MMR activity by negatively regulating the PMS2 or MLH1-PMS2 level on chromatin. Nevertheless, we noticed that overexpression of MRE11A greatly decreased chromatin bound PMS2 protein level while only mildly, though statistically significantly, downregulated MMR activity, which may imply the possible backup role of MRE11A as an endonuclease, though not as efficient as PMS2, in MMR pathway.

In conclusion, our study identified MRE11A as one novel negative regulator of MMR activity. Furthermore, based on our results and previous studies, we propose that the binding of MRE11A to MLH1 sterically hinders the binding of PMS2 to MLH1, hence downregulating the MLH1-PMS2 heterodimers on chromatin **(Figure 7)**. We speculate that the competition between PMS2 and MRE11A for binding to MLH1 is a regulatory circuit to ensure balanced MMR activity. Further investigations may focus on elucidating the underlying biological importance of the interaction between MRE11A and MLH1 in the maintenance of genomic stability not only through balancing the MMR activity but also in alternative DNA metabolism pathways, such as double strand breaks repair and processing of replication stress.

**Figure 7.**
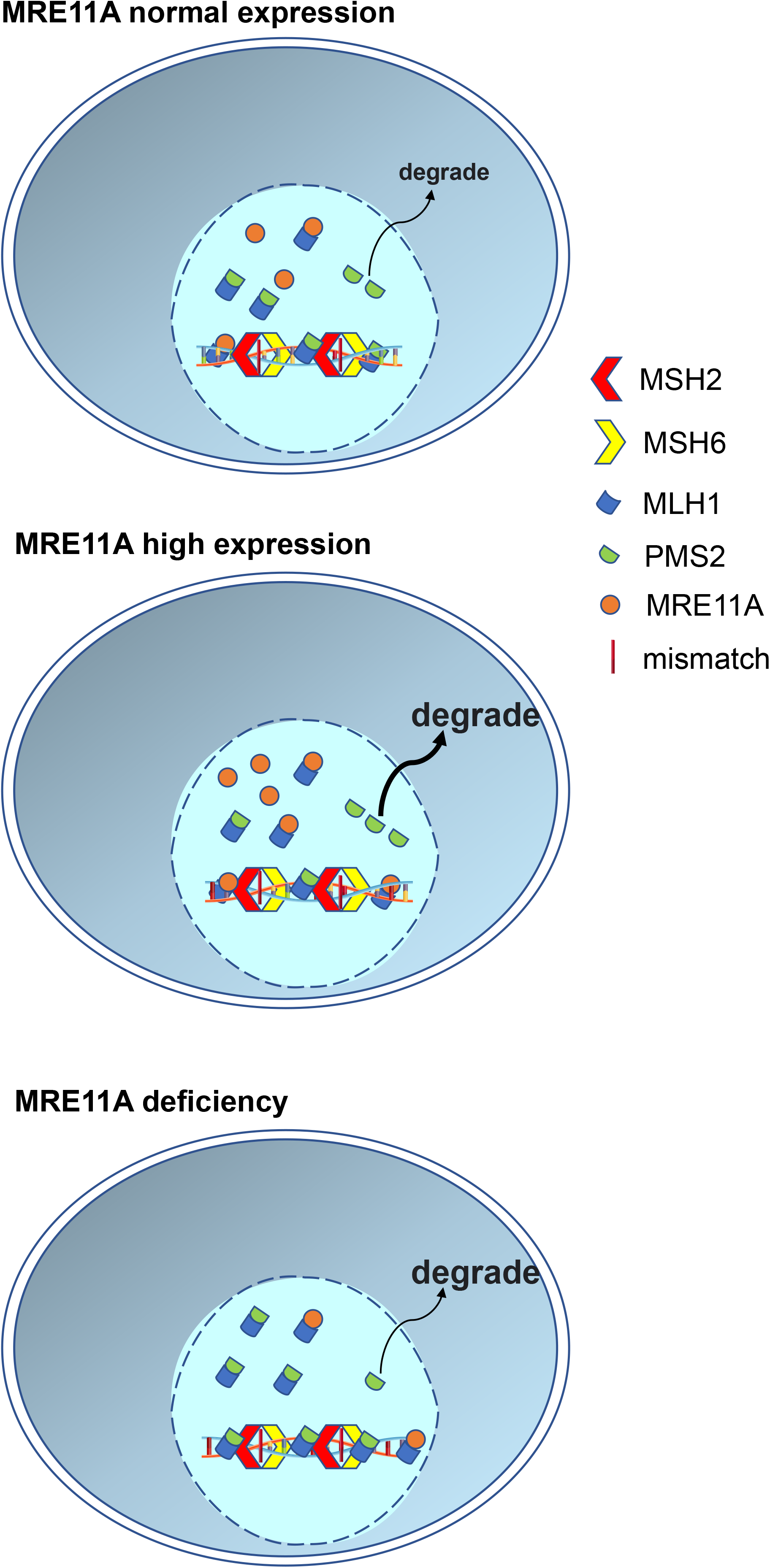
Schematic summary of the study. In naïve cells, a proportion of MRE11A may interact with MLH1, but do not interfere the proper interaction between intrinsic PMS2 and MLH1. In MRE11A overexpression cells, excessive MRE11A occupied the binding site of PMS2 to MLH1, leading to the degradation of unbound PMS2 and decreased MLH1·PMS2 heterodimer on chromatin, consequently compromising the MMR activity. While in MRE11A deficiency cells, more intrinsic PMS2 bind to MLH1, leading to increased MMR activity and thus more sensitive to MNNG treatment.

## MATERIAL AND METHODS

### Cell lines generation

Hela cells were purchased from CELLCOOK and STR identified in advance. All cells were grown in DMEM (Bioind) containing 10% Fetal Bovine Serum (Bioind) and 1% penicillin/streptomycin (Gibco) at 37°C in 5% CO2. The MRE11 knock-down cell lines were generated by lentivirus obtained from GeneChem. Two different target sequences were respectively used (siMRE11-1: 5′-GTACGTCGTTTCAGAGAAA-3′, siMRE11-2: 5′-GGAGGATATTGTTCTAGCT-3′).

### Gene overexpression and knockdown

The MRE11A (NM_001330347) and PMS2 (NM_000535) overexpression constructs were obtained through gene synthesis and cloned into pcDNA3.1(+) by Gene Universal.

Flag-MRE11A and Flag-MRE11A (452-634 aa) cassettes were achieved by PCR using the following primers: Fwd 5′-GGGGTACCgccaccatgGATTACAAGGACGACGATGACAAGagtactgcagatgcact-3′ and Rev 5′-AATGCGGCCGC TTATCTTCTATTTCTTCTTAAAG-3′ for Flag-MRE11 and Fwd 5′-GGGGTACCgccaccatgGATTACAAGGACGACGATGACAAGagagggatgggtgaagcagt 3′-and Rev 5′-AATGCGGCCGCTTAATTTCGGGAAGGCTGCTGTC-3′ for Flag-MRE11 (452-634 aa).

The PCR products were then subcloned into pcDNA3.1(+) plasmid. For each 6-well plate, 2 μg plasmid were transfected using PolyJet™ In Vitro DNA Transfection Reagent (SignaGen, SL100688) according to the manufacturer’s instructions.

All siRNAs were synthesized by RiboBio according to the following target sequences: siMRE11A-1: 5′-GTACGTCGTTTCAGAGAAA-3′, siMRE11-2: 5′-GGAGGATATTGTTCTAGCT-3′, siMSH2: 5′-GCTAAAAGCTGAAGTAATA-3′, siMLH1: 5′-CTGAGATGCTTGCAGACTA-3′, siNC (negative control): 5′-TTCTCCGAACGTGTCACGT-3′. siRNA transfection was followed the general guidelines of reverse transfection using Lipofectamine® 2000 (Thermo Scientific, 11668019). In brief, 5μl Lipofectamine® 2000 and 100 pm siRNA were respectively diluted in 200 μL Opti-MEM™ I Reduced Serum Medium (Thermo Scientific, 31985062) for 5 min at RT, then added the diluted Lipofectamine® 2000 and siRNA to the 6-well plate, mix gently and incubate for 15 minutes at room temperature. After that, 2ml complete growth medium without antibiotics with 1×10^6^ cells were added to the plate and mix gently. Incubated the cells at 37°C with 5% CO2 and harvested after 96 hours.

### RNA extraction and RT-qPCR analyses

Total RNA Isolation Kit was purchased from Vazyme (RC101). For RT-qPCR, RNA was reverse transcribed to cDNA by Reverse Transcriptase (Vazyme, R312-01). qPCR analyses were performed with Universal SYBR qPCR Master Mix (Vazyme, Q711-02). For the results analyze, GAPDH was used as reference. The primers were listed below: MRE11A: Fwd 5′-ATCGGCCTGTCCAGTTTGAAA-3′ and Rev 5′-TGCCATCTTGATAGTTCACCCAT-3′. PMS2: Fwd 5′-TTTGCCGACCTAACTCAGGTT-3′ and Rev 5′-CGATGCGTGGCAGGTAGAA-3′. GAPDH: Fwd 5′- GGAGCGAGATCCCTCCAAAAT -3′ and Rev 5′-GGCTGTTGTCATACTTCTCATGG -3′

### Western blotting and antibodies

Cell lysis solution were purchased from Beyotime (P0013B), containing 50mM Tris(pH 7.4),150mM NaCl, 1% Triton X-100, 1% sodium deoxycholate and 0.1% SDS, then added Phosphatase inhibitor (Beyotime, P1081) and Protease Inhibitor (Cwbio, CW2200) just before use. For protein extraction, aspirated media from plates, washed cells thrice with cold PBS, then added lysis solution to the plate and placed on ice for 30 min. Immediately collected the lysis solution in a microcentrifuge tube and centrifuged at 12000 rpm for 5 min (4°C), and supernatants were collected as the total cell extracts. Protein levels were determined using BCA Protein Assay Kit (Beyotime, P0012). For western blot, after SDS-PAGE gels electrophoresis, proteins were transferred to PVDF membrane and blocked by TBST (137 mM NaCl, 20 mM Tris, 0.1% Tween 20) with 5% (w/v) skim milk for 1 hour at room temperature. Next, incubated membrane with primary antibody in antibody dilution buffer (TBST containing 5% skim milk) overnight at 4°C. The next day, the membrane was washed three times for 5 min each with TBST, and incubated with the secondary antibody for 1 hour at room temperature. Protein signals were visualized with ECL Western Blotting Reagents (Vazyme, E412-02). Primary antibodies are as followings: Flag (Proteintech, 20543-1-AP), PMS2 (Proteintech, 66075-1-Ig), MSH6 (Proteintech, 18120-1-AP), EXO1 (Proteintech, 16253-1-AP), Histone-H3 (Proteintech, 17168-1-AP), CHK2 (Proteintech, 13954-1-AP), CHK1 (Proteintech, 25887-1-AP), MSH2 (Proteintech, 15520-1-AP) MRE11A (Proteintech, 10744-1-AP), actin (Proteintech, 20536-1-AP), P-CHK2-Thr68 (CST, #2661), P-CHK1-Ser345 (CST, #2348); MLH1 (Affinity, DF6057). Secondary Antibodies: HRP-conjugated Affinipure Goat Anti-Rabbit IgG (SA00001-2) and HRP-conjugated Affinipure Goat Anti-Mouse IgG (SA00001-1), were purchased from Proteintech.

### Chromatin extraction and co-immunoprecipitation

Chromatin extraction was performed with the Chromatin Extraction Kit (Abcam, #ab117152) according to the manufacturer instructions. Briefly, cells were trypsinized and washed twice with 10 mL cold PBS and count the cells with hemacytometer. Then harvest the cell pellet by centrifuge. Next, Lysis Buffer (200 µL/ 1×10^6^ cells) contained Protease Inhibitor was added to the cell pellet and resuspended gently, incubated on ice for 10 min and vortex vigorously for 10 sec, removed supernatant by centrifuge. The sediment was resuspended using Extraction Buffer (50 µL/1×10^6^ cells) contained Protease Inhibitor and incubated the on ice for 10 min and vortex occasionally, followed by sonication and centrifuged at 12,000 rpm at 4°C for 10 min and supernatants were collected as chromatin extracts. The proteins bonded with chromatin were analyzed with western blotting.

For immunoprecipitation, extracted chromatin proteins were incubated with MSH2 antibody (Proteintech, 15520-1-AP) or normal rabbit IgG (CST, #2729) overnight at 4°C and then with Protein G Magnetic Beads (MCE, HY-K0204) for 1 hour at RT followed by washed three times with PBST (1×PBS with 0.5% Tween-20, pH 7.4). Then Beads were separated using Magnetic Separation Rack, and the beads were washed three times with PBST again. Then 1×SDS-PAGE Loading Buffer was added and heated sample to 98°C for 5 min. Finally separated the Beads by centrifuge and transferred supernatant containing proteins to a new vial. The products were analyzed by western blotting.

### Immunofluorescence

The cells were plated on 6-well Cell Culture Plate with coverslips. To assess 53BP1 foci, cells were treated with O6-Benzylguanine (sigma) for 1 hour, then treated with MNNG for 12 hours. Following MNNG treatment, cells were fixed with 4% paraformaldehyde for 10 min, permeabilized in PBS buffer containing 0.1% Triton X-100 for 15 min. After that, cells were incubated with 53BP1 antibody (Cell signaling, #4937) and CYCLIN A antibody (santa, sc-271645) overnight at 4°C and then stained with Alexa Fluor 488 or Alexa Fluor 555 secondary antibodies (Invitrogen) for 1 hour at RT. Nuclei were counterstained with DAPI. Finally, coverslips were mounted with anti-fading (Solarbio). Fluorescent images were taken by Leica DM6 B fluorescence microscope.

### Cell survival, growth and apoptosis analysis

N-Methyl-N′-nitro-N-nitrosoguanidine (MNNG) and O6-Benzylguanine (O6-BG) were dissolved in DMSO and stored at -20°C. Hela cells were treated with O6-BG (10 μM) for 1 hour prior to addition of MNNG to at the indicated concentration.

To assess the survival, cells were washed and harvested after treatment with indicated concentrations of MNNG for 72 hours and resuspended in 1ml PBS, then measured the optical density (OD) at 600 nm using an iMark™ Microplate Absorbance Reader (Bio-Rad).

For clongenic assay, Hela cells were transfected with scrambled or targeted siRNA and cells were plated on 6-well plates one day before treated with indicated concentrations of MNNG. Approx. 8 days after treatment, cells were stained with 0.5% crystal violet in 20% ethanol. Only colonies containing >100 cells were counted.

To measure the growth rate, suspended cells were counted using hemacytometer and diluted to 10000 cells per milliliter, then inoculated into a 96-well plate (100μL per well) in sextuplicate. After 24, 48, 72 and 96 hours of incubation at 37°C, 10μL of CCK-8 (Vazyme, A311-01) was added to each well for 1 hour incubation. After that, plate was measured the absorbance (OD) at 450 nm by the iMark™ Microplate Absorbance Reader (Bio-Rad). The growth rate was defined as the OD ratio to the OD at 24 hours.

To detect apoptosis, cells were treated with MNNG, and harvested after 72 hours, washed twice and suspended in binding buffer. Then, stained the cells by a Annexin V-FITC/PI Apoptosis Detection Kit (Vazyme, A211-01) and detected cells distribution using a Beckman Coulter Gallios (Beckman Coulter). The result was analyzed by Kaluza Analysis 2.1 (Beckman Coulter).

### Microsatellite Analysis and MMR assay

We isolated single cell using the limiting dilution approach, and grew for approx. 30 generations to permit mutation accumulation. We then collected ten clones from each treated group and sent to Shanghai Personalbio Technology Co., Ltd. for MSI analysis based on the fluorescent PCR amplification of microsatellite genes, including NR-21, NR-24, BAT-25, BAT-26, MONO-27, and capillary electrophoresis analysis by 3730xl DNA Analyzer. The results were demonstrated by GeneMaker software.

For MMR assay in cells, substrate heteroduplex GFP substrate was prepared according to Zhou et al. (Zhou *et al*., 2009). Briefly, p111 was nicked with Nb.Bpu10I (Thermo Scientific) and further digested with ExoIII (NEB) to generate single strand circular DNA. p189 was linearized by SphI (NEB). To obtain the GFP-heteroduplex, the linearized double strand DNA was annealed with p111 single strand circular DNA in 1 X annealing buffer (Beyotime, #D0251) by heated at 95°C for 5min and followed by slowly cooling down from 95°C to 25°C within 45 min. The annealing product was then treated by Plasmid-Safe ATP-Dependent DNase (Biosearch Technologies). To assess MMR activity, Hela cells were transfected with 1μg of the heteroduplex plasmid and 0.8μg of pmCherry-C1. After incubation for 24 hours the cells were harvested and analyzed for fluorescence intensity with a Beckman Coulter Gallios (Beckman Coulter). The result was analyzed by Kaluza Analysis 2.1 (Beckman Coulter) and relative repair efficiency was measured by the ratio of GFP-positive cells to that of mCherry-positive cells.

### Statistical analyses

All data were presented as the mean ± SD and analyzed by unpaired t-tests.. The light and fluorescence microscopy photos were captured by Leica LAS X software. ImageJ were used for intensity quantifications of blotting assays and the FACS data were analyzed and illustrated by Kaluza Analysis Software. Statistical analyses were performed by GraphPad Prism 8. Statistical significance was set when *P* < 0.05 (two sided).

## ACKNOWLEDGEMENT

We thank Dr. Lu-Zhe Sun for the kindly donation of P111 and P189 plasmids. This work is supported by National Natural Science Foundation of China [grant number 31800682, 82211540400] and Safety Evaluation of Chinese Materia of Nanjing University of Chinese Medicine [grant number JKLPSE201814].

## AUTHOR CONTRIBUTION

Du D and Yang Y have equal contribution to the manuscript. Du D, Liu D and Yang Y performed experiments. Wang G and Chen L assisted the experiments. Guan X and Liu D analyzed the data and organized the figures. Lene J assisted to analyze data and suggested the idea. Guan X edited the manuscript and supervised the study. Liu D wrote the manuscript and developed the overall concept.

## CONFLICT OF INTEREST

The authors have no conflict of interests to declare.

## Data Availability Section

This study includes no data deposited in external repositories.

## Supplementary figure legends

**Figure S1.**
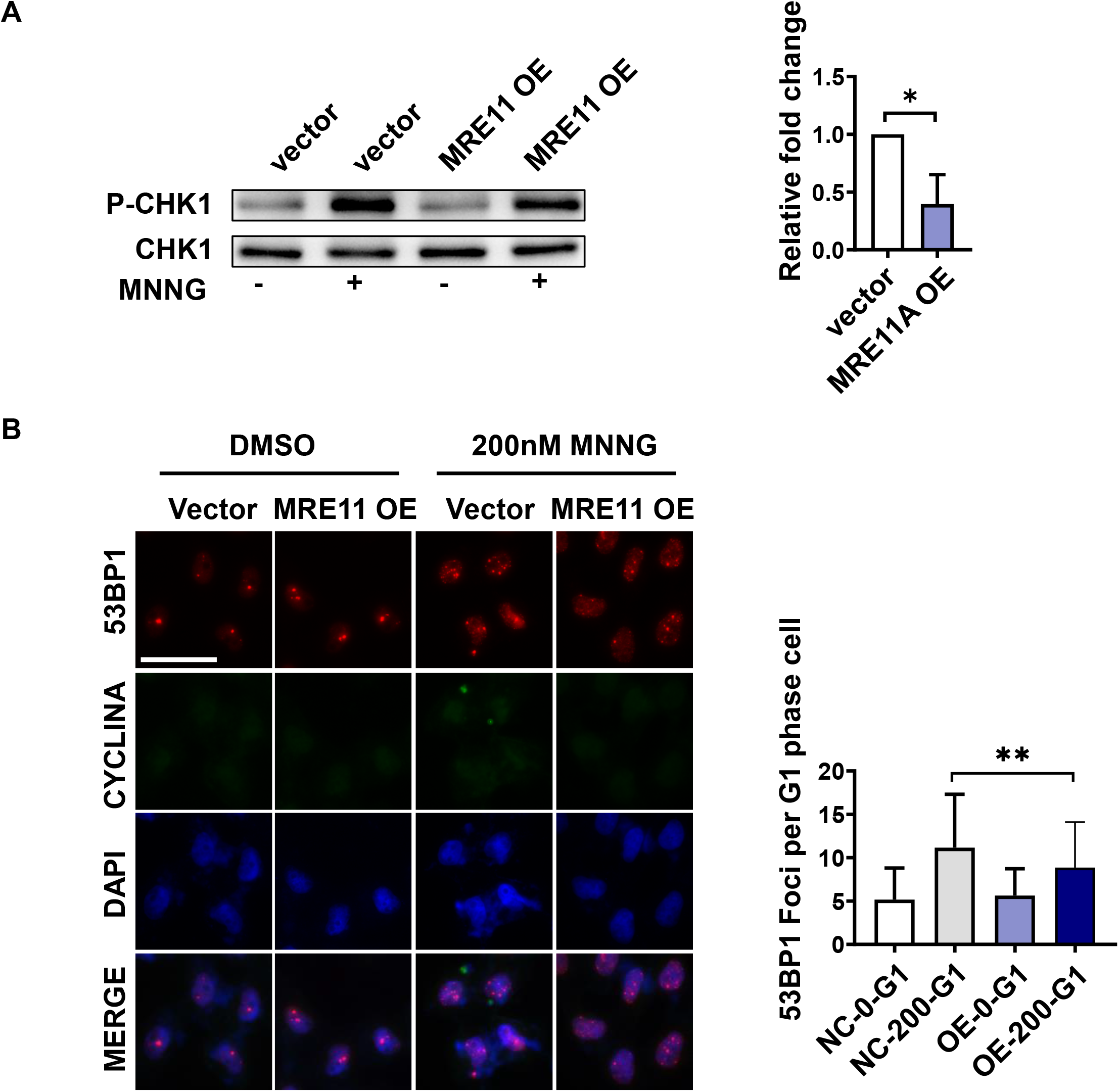
MRE11A overexpression decreases levels of DNA damage signals 12h after MNNG treatment. **(A)** The representative western blotting of the phosphorylation levels of CHECK1 12 hours after DMSO or 200nM MNNG treatment. The alternation of phosphorylation level was calculated as the p-CHECK1 level to total CHECK1 protein after 200nM MNNG treatment minus that with only DMSO treatment. Right graph showed the quantification of proteins level changes relative to siNC. **(B)** Representative Immunofluorescent pictures of the 53BP1 foci in G1 phase 12 hours after DMSO or 200nM MNNG treatment. Right graph showed the quantification of the number of 53BP1 foci per cell in G1 phase (CYCLINA+). Data are shown as mean ± SD, n = 3, * p < 0.05, ** p < 0.01, *** p < 0.001, using unpaired two-tailed Student’s t test.

**Figure S2.**
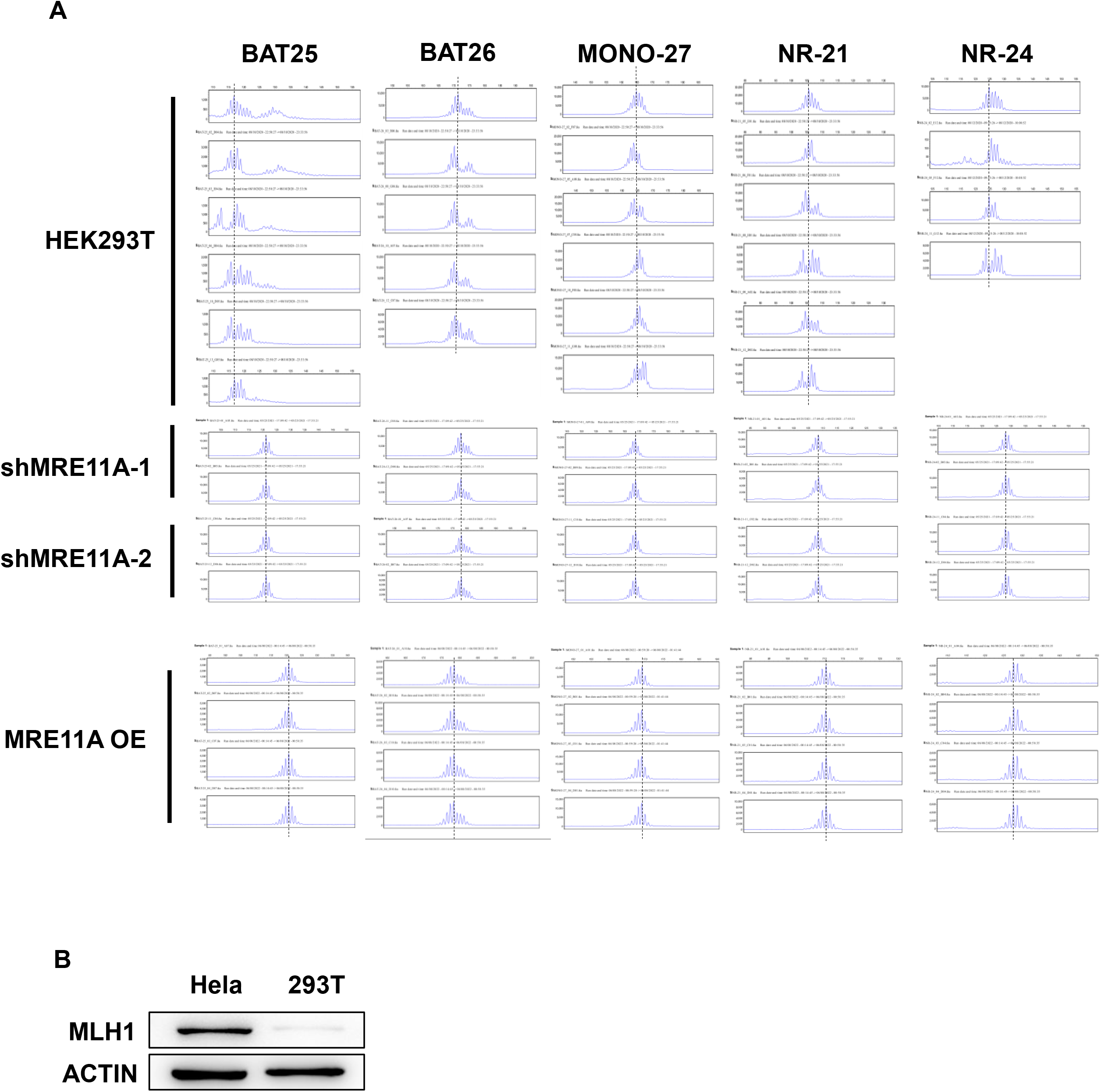
MRE11A alternations does not induce microsatellite instability. **(A)** Capillary electrophoresis of PCR amplification products of indicated microsatellite gene loci was used for MSI test. Each group included ten samples and 293T cell was set as positive control and only representative analysis results were shown. **(B)** Western blotting results of MLH1 levels in Hela cells and 293T cells.

